## Supplementary Information for "Lipid packing and cholesterol content regulate membrane wetting and remodeling by biomolecular condensates"

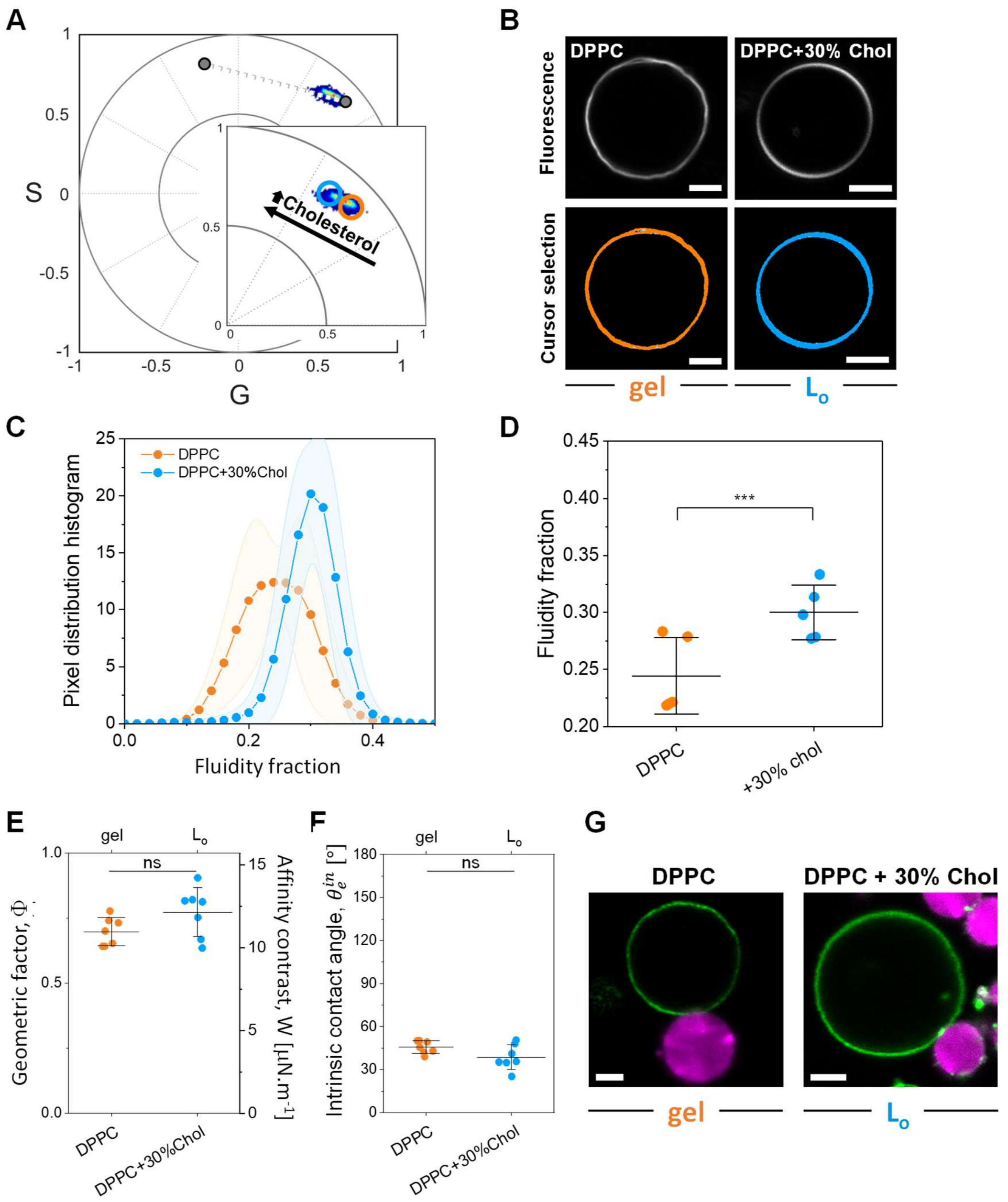


**Figure S1.** (**A**) Spectral phasor plot for LAURDAN in GUVs composed of DPPC or DPPC:Chol 7:3 at (23±1)°C, in the gel ($L_{\beta'}$) or the liquid ordered phase ($L_o$), respectively. The phasor position shifts counterclockwise, indicating a more fluid environment when cholesterol is present in the membrane, as expected. Note that the trajectory described by the data aligns with the one shown in Fig. 1B and 3A. (**B**) Representative confocal images of DPPC and DPPC:Chol 7:3 GUVs labeled with 0.5 mol% LAURDAN. The images in the bottom panel are painted according to the circular cursors shown in (A). (**C**) Pixel distribution histogram along the linear trajectory drawn as a white dotted line in (A), showing the fluidity fraction for the different membrane compositions. Data are represented as the mean (circles and lines) ± SD (shadowed contour), n = 5 independent experiments per condition. (**D**) Center of mass of the histograms shown in (B). (**E**) Geometric factor, Φ, and affinity contrast, W, for DPPC and DPPC:Chol 7:3 GUVs in contact with glycinin condensates in 150 mM NaCl. Individual data points are shown for each membrane composition. The lines indicate the mean value ± SD. (**F**) Intrinsic contact angle, $\theta_e^{in}$ for the systems in (E). Individual data points are shown for each membrane composition. The lines indicate the mean value ± SD. (**G**) Representative confocal images of DPPC and DPPC:Chol 7:3 GUVs in contact with glycinin condensates at 150 mM NaCl. All scale bars: 5 μm.

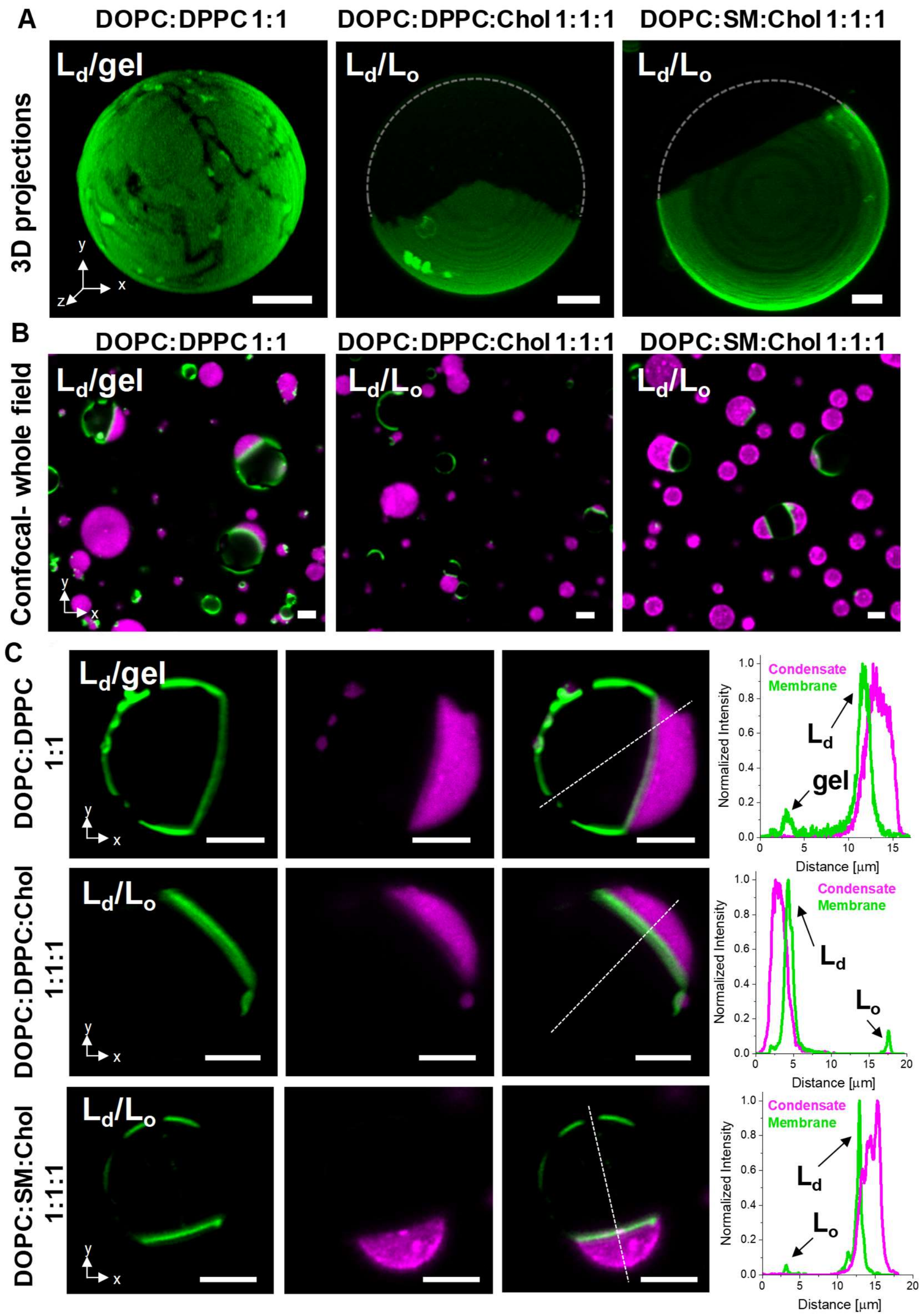


**Figure S2.** (**A**) 3D projections of vesicles composed of DOPC:DPPC 1:1, DOPC:DPPC:Chol 1:1:1, and DOPC: SM:Chol 1:1:1 labeled with 0.1 mol% ATTO 647N-DOPE (green). Because ATTO 647N-DOPE preferentially partitions into the liquid-disordered phase, fluorescence is brighter in this phase, while the gel and liquid-ordered phases appear darker. (**B**) Large field image showing vesicles of the indicated compositions in contact with FITC-labeled glycinin condensates (magenta). In all cases the condensates only interact with the phase presenting lower lipid packing ($L_d$). (**C**) Examples of vesicles of the indicated binary and ternary mixtures in contact with glycinin condensates. Individual channels are shown with the corresponding line profiles indicating that condensates only interact with the less packed phases (liquid-disordered). All scale bars: 5 µm.

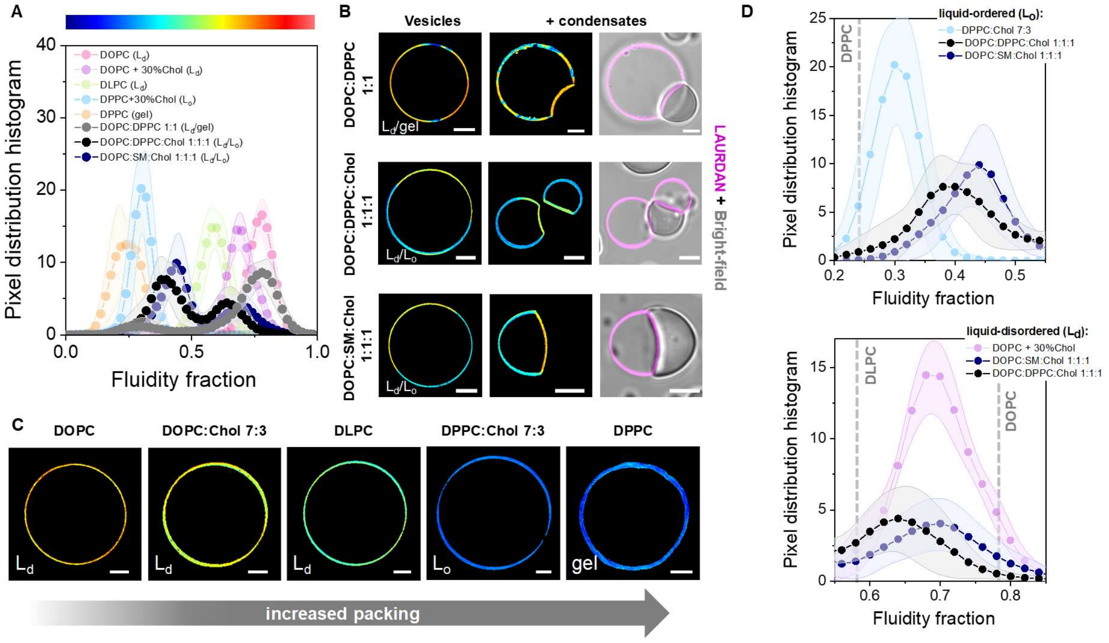


**Figure S3. (A)** Pixel distribution histogram vs fluidity fraction showing an example for DOPC:DPPC 1:1 (gray circles), DOPC:DPPC:Chol 1:1:1 (black circles), and DOPC:SM:Chol (blue circles) vesicles labeled with 0.5 mol% LAURDAN. The fluidity fractions for the compositions in Figures 1, 3 and S1 are included here for comparison. (**B**) A continuous color scheme (rainbow) is assigned to the fluidity fraction, as shown on top of panel (A), and the images for DOPC:DPPC 1:1, DOPC:DPPC:Chol 1:1:1, and DOPC:SM:Chol without and in contact with condensates are colored accordingly. As the condensates are not labeled, bright-field images merged with the LAURDAN channel are included for reference. (**C**) Images of vesicles of the indicated compositions colored with the continuous color scheme shown in (A). (**D**) Zoomed panels of the plot shown in (A), highlighting the differences between the various liquid-ordered ($L_o$) phases (upper panel), and liquid-disordered ($L_d$) phases (lower panel) for the binary and ternary mixtures. The maximum position for the single lipid compositions is indicated with gray dashed lines for reference. The fluidity for the liquid-ordered phase increases in the order: DOPC:Chol 7:3 < DOPC:DPPC:Chol 1:1:1 < DOPC:SM:Chol 1:1:1:, while for the liquid-disordered phase the fluidity increases according to: DOPC:DPPC:Chol 1:1:1 < DOPC:Chol 7:3 ≤ DOPC:SM:Chol 1:1:1. All scale bars: 5 µm.

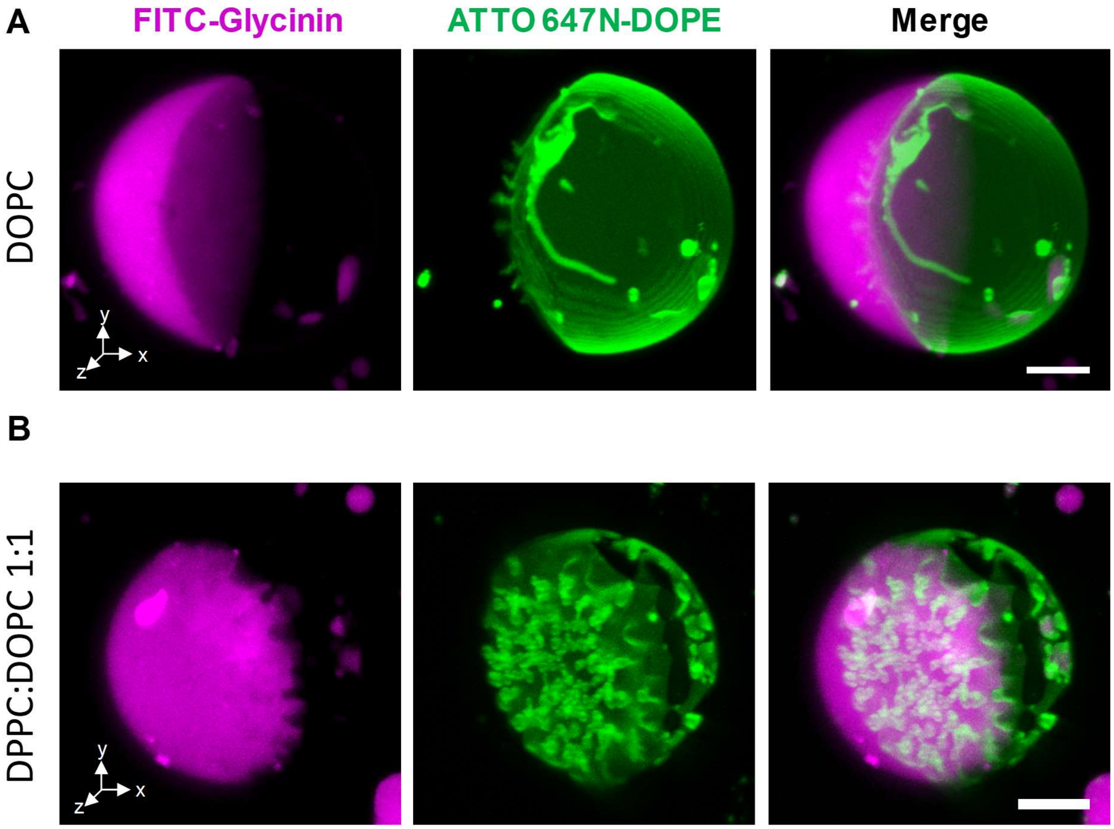


**Figure S4.** Confocal 3D projections of the vesicles shown in Fig.5: (**A**) DOPC and (**B**) DPPC:DOPC 1:1 GUVs (green) in contact with glycinin condensates (magenta) displaying tubulation of the interfacial region. Scale bars: 5 µm.

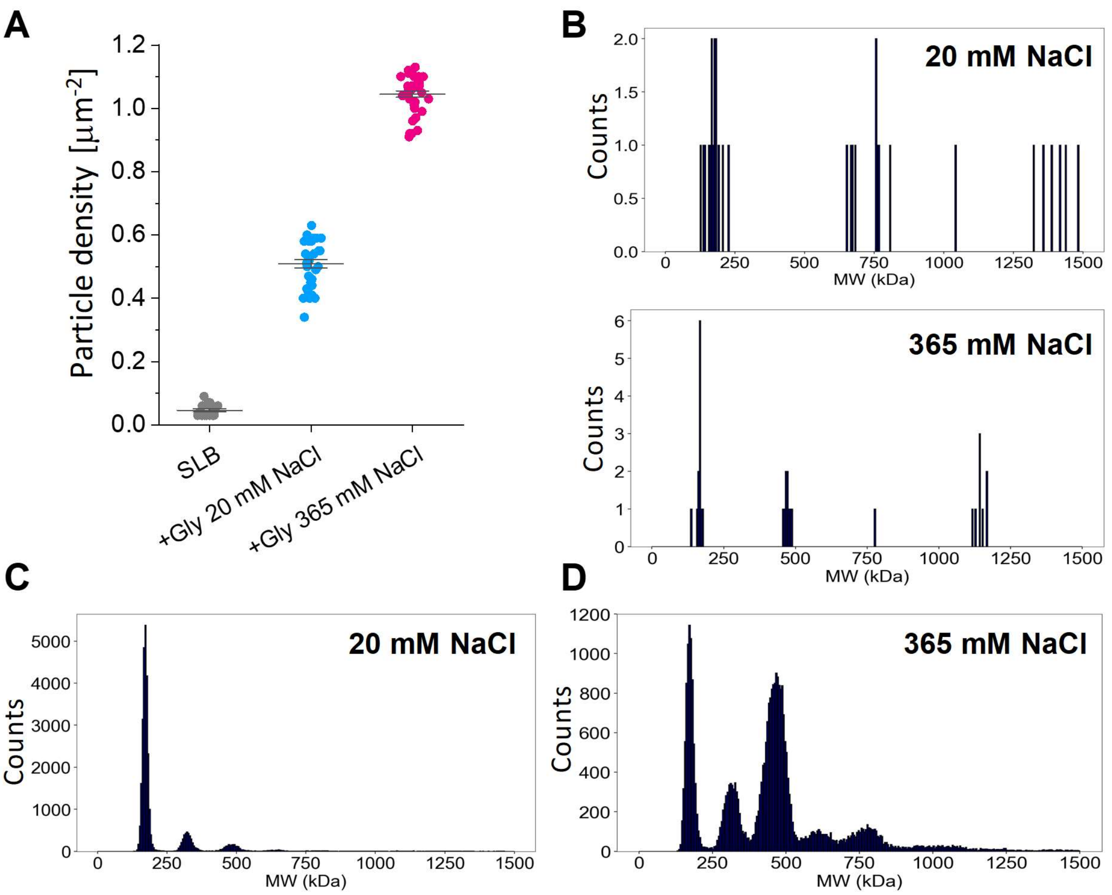


**Figure S5.** Mass photometry data. (**A**) Particle density obtained for DOPC supported lipid bilayers (SLB) in the absence of protein or in contact with 0.48 µg/mL glycinin solutions at the indicated NaCl concentrations (same data as in Figure 6F, here showing also the background signal of the SLB). (**B**) Particle counts vs molecular weight (MW) for DOPC SLB in the absence of protein at the indicated NaCl concentrations assessed from N = 7 (20 mM), and N = 9 (365 mM NaCl) recorded movies. The detected particles arise from impurities or unfused vesicles. (**C**) Protein mass distribution plot for DOPC SLB at 20 mM NaCl, and (**D**) at 365 mM NaCl (note the much higher particle count compared to the data in panel B). At this protein concentration, glycinin predominantly adsorbs as trimer (160 kDa) at 20 mM NaCl, and additionally as hexamer (320 kDa) formed by two trimer subunits[1]; and higher oligomeric species. At high salinity, glycinin forms more of the higher oligomeric species. The depicted plots in (C) and (D) represent combined data from N = 31, and N = 34 recorded movies, respectively. The mass distribution plots in (B-D) show the mean particle mass of each particle detected over a time period of at least 50 frames (186 ms).

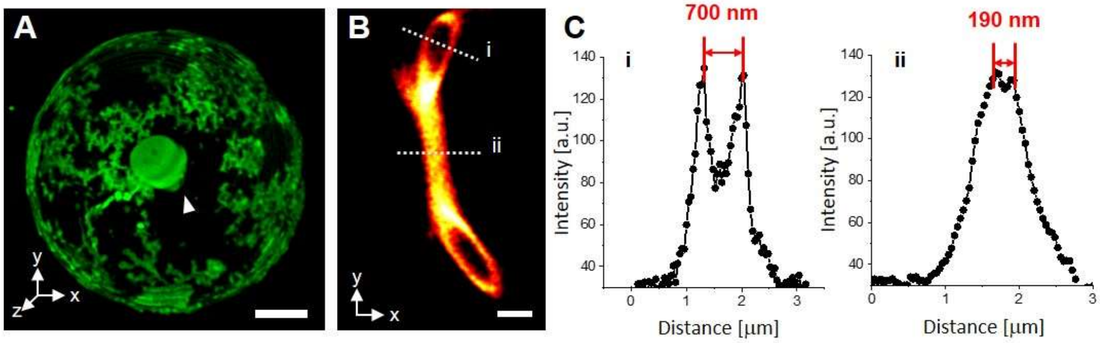


**Figure S6.** (**A**) Confocal 3D projections of a DOPC GUV labeled with 0.1 mol% ATTO 647N-DOPE in contact with a homogeneous solution of glycinin (10 mg/mL in 365 mM NaCl) displaying nanotube and membrane sheet formation. (**B**) STED image of a membrane sheet attached to a vesicle. (**C**) Intensity profiles of the dashed lines indicated in (B) providing information about the thickness at the center and periphery of the cisterna-like double membrane sheet. Scale bars: 5 µm.

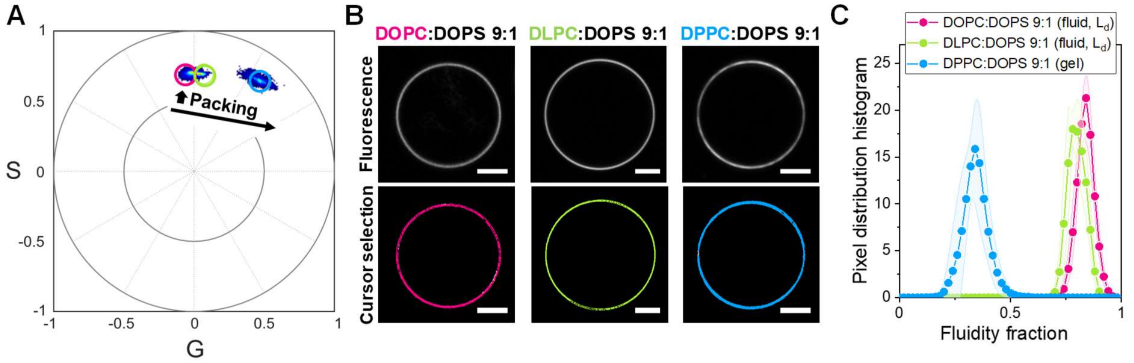


**Figure S7.** (**A**) LAURDAN spectral phasor for vesicles composed of DOPC, DLPC, and DPPC containing 10 mol% of DOPS. (**B**) Representative confocal images for the different compositions (upper panel) and cursor painted images (lower panel) according to the pixels selected in (A). Scale bars: 5 µm. (**C**) Pixel distribution histogram of the fluidity fraction for DOPC, DLPC, and DPPC membranes containing 10 mol% of DOPS.

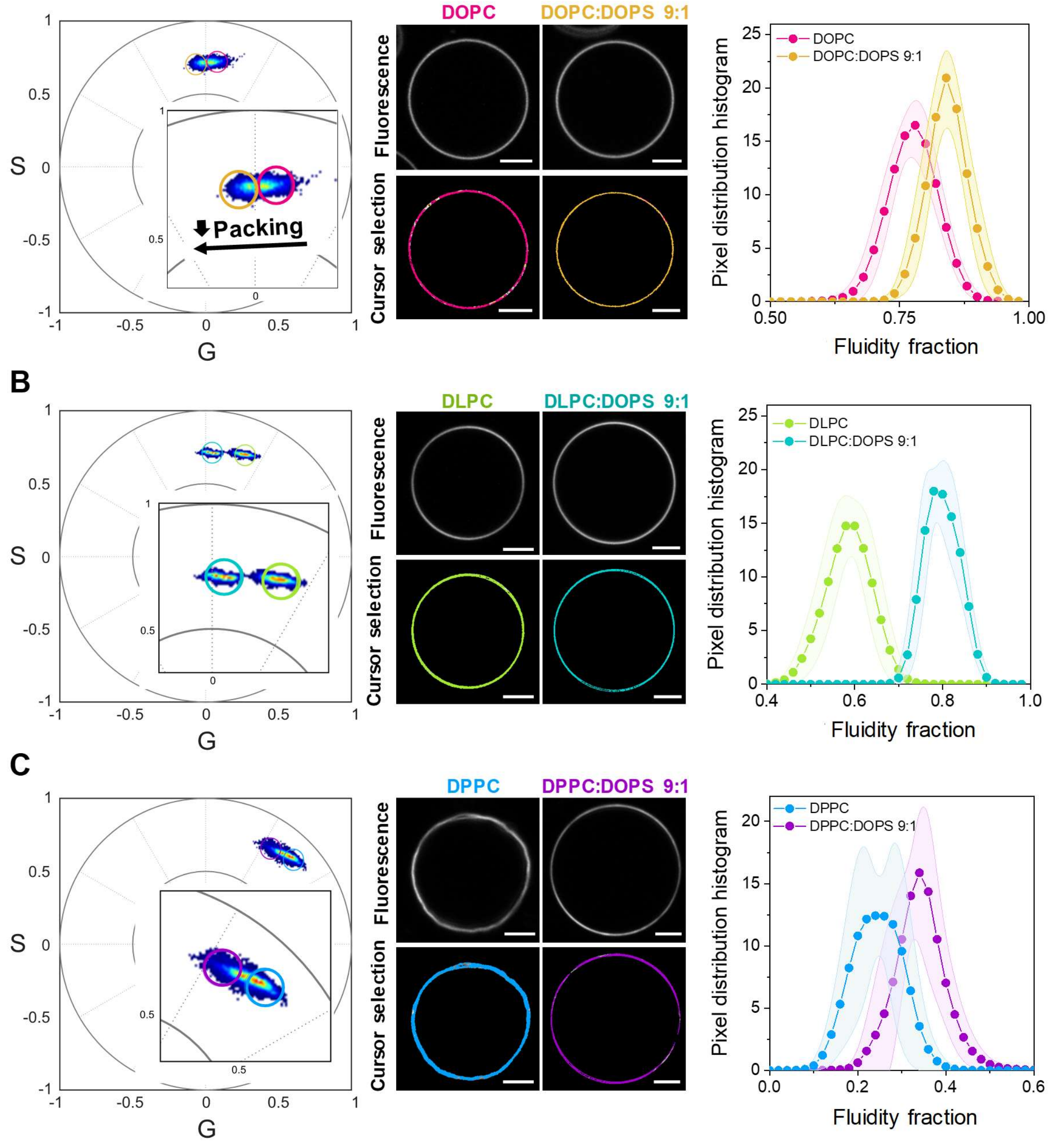


**Figure S8.** LAURDAN spectral phasors, representative confocal and cursor painted images, and pixel distribution histograms for GUVs composed of DOPC:DOPS 9:1 (**A**), DLPC:DOPS 9:1 (**B**), and DPPC:DOPS 9:1 (**C**) and the respective DOPS-free membranes. Note that in all cases the addition of the charged DOPS decreases the lipid packing (i.e. increases the fluidity fraction). Scale bars: 5 μm.

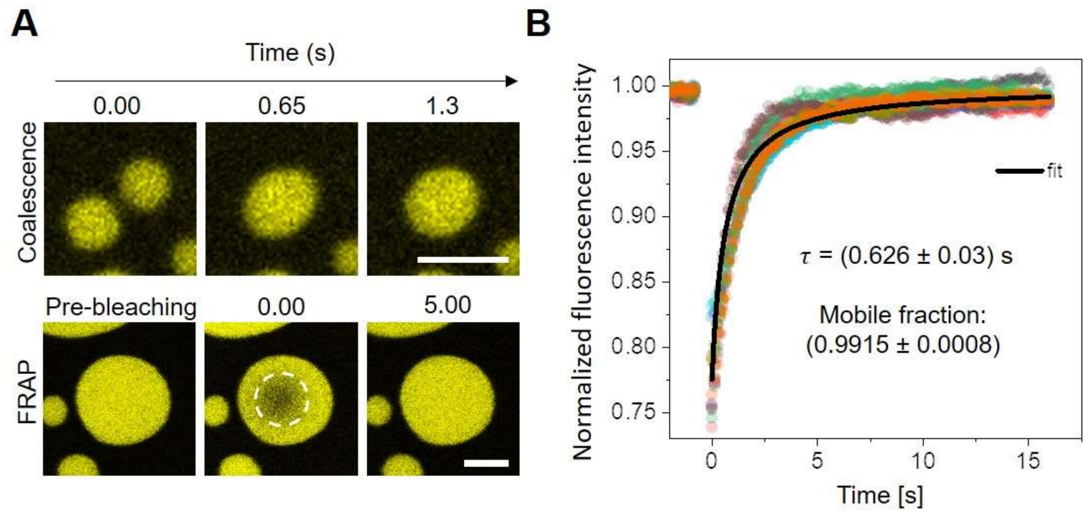


**Figure S9.** (**A**) Upper panel: $K_{10}/D_{10}$ condensates coalescence; lower panel: TAMRA-$K_{10}$ fluorescence recovery after photo-bleaching (FRAP) in $K_{10}/D_{10}$ condensates. Scale bars: 5 µm. (**B**) FRAP curves for TAMRA-$K_{10}$ in $K_{10}/D_{10}$ condensates. 10 independent measurements are shown, and the fitting corresponds to the function: $y=(I_0+I_{max}(x/\tau_{1/2}))/(1+x/\tau_{1/2})$, where $I_0$ is the initial intensity, $I_{max}$ is the maximal intensity, and $\tau_{1/2}$ is the halftime of recovery. The apparent diffusion coefficient was calculated using the relation: $D_{app}=r_0^2\ \upsilon/4\ \tau_{1/2}$, where $r_0$ is the radius of the bleaching spot and $\upsilon$ is a correction factor accounting for the difference between the defined size of bleaching spot and its real size. Using the diffusion coefficient, the apparent viscosity ($\eta$) was estimated from the Stokes-Einstein relation: $\eta_{app}=k_BT/6\pi R_hD_{app}$, where $R_h$ is the hydrodynamic radius estimated as $R_h$=1.02 nm for $K_{10}$[2]. The obtained apparent viscosity is $\eta_{app}$=80 mPa·s, which is within the order of magnitude of previously reported data for this system[2]. As the speed of condensate coalescence was faster than our confocal setup (see upper panel in A), it was not possible to have an estimation of the inverse capillary velocity ($\eta/\Sigma_{ce}$) to derive the interfacial tension ($\Sigma_{ce}$). We assume that the interfacial tension should be similar to that reported for polyK systems, $\Sigma_{ce}$=17µN/m[3].

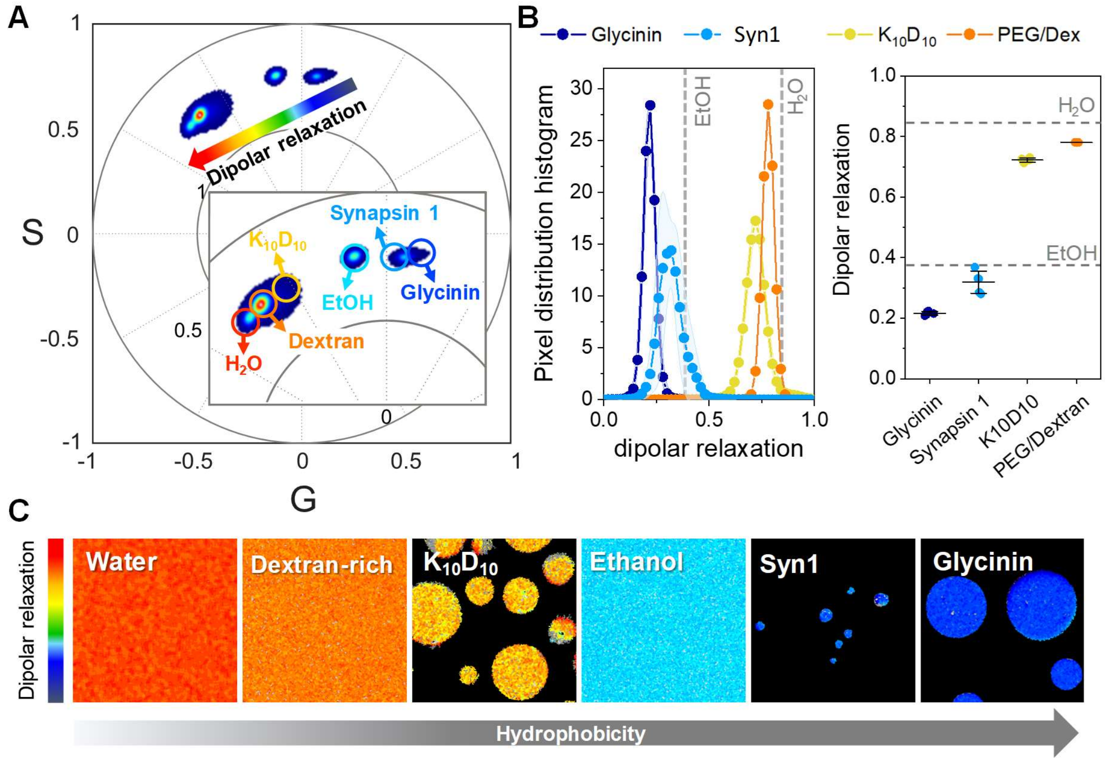


**Figure S10.** Condensates micropolarity measured by ACDAN spectral phasors. (A) Spectral phasor plot of ACDAN in the various condensate systems, with reference data for ACDAN in water and ethanol (EtOH) included. For PEG/dextran condensates, measurements were conducted directly in the bulk dextran-rich phase of the phase-separated system. (B) Left panel: pixel distribution histograms of the data shown in (A). Right panel: center of mass of the distributions plotted in the left panel highlighting differences in micropolarity across the condensate systems. The values for water and EtOH are indicated with dashed lines for reference. (C) Cursor-colored images illustrating variations in dipolar relaxation between the points indicated by the arrow in (A) providing direct visualization of micropolarity differences.

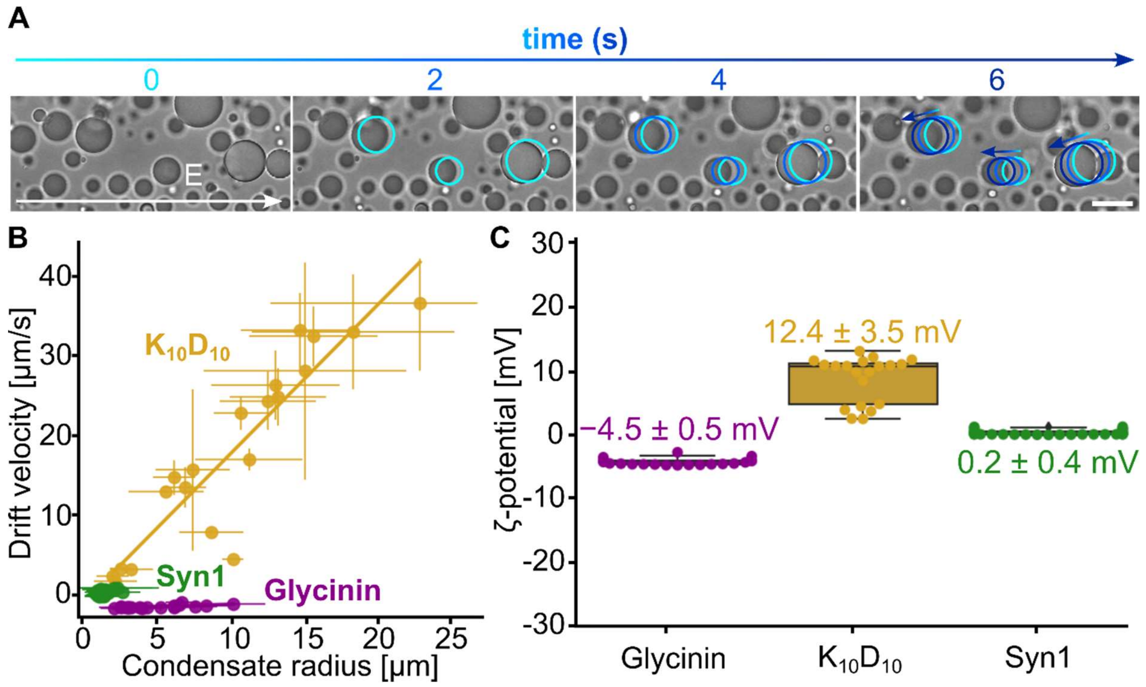


**Figure S11.** Condensates ζ-potential measured by the microelectrophoresis method[4] (see Material and Methods for details). (A) Bright-field microscopy images showing the time sequence of the displacement of glycinin condensates in the opposite direction of the externally applied electric field (E=5 V/cm). The colored circles are guides to the eye highlighting the trajectory of condensates positions. Scale bar is 20 μm. (B) Drift velocity of condensates with different radius migrating in electric fields of 5 V/cm (glycinin and $K_{10}D_{10}$) and 10-30 V/cm (Syn1) (C) ζ-potentials of the different condensates computed from equation (7). Individual data points are shown, and the mean±SD values are indicated on top of each box plot.

**Table S1: Summary of the material properties of the tested condensate systems.**

| Condensate | Viscosity (Pa.s) | Surface tension (μN/m) |
|---|---|---|
| **Glycinin** (ref. 5) | 195 | 15.7 |
| **Synapsin 1** (ref. 6) | 250 | 23 |
| **PEG/dextran** (ref. 7 and 8) | 0.07 | 8 |
| **$K_{10}/D_{10}$** (ref. 3 and this work) | 0.08 | 17 |

**Supplementary movies:**

**Movie S1:** Confocal microscopy z-stack of a giant vesicle composed of DOPC:DPPC 1:1 labeled with 0.1 mol% ATTO 647N-DOPE (green) displaying fluid/gel phase coexistence in contact with a glycinin condensate containing 4 %v/v of FITC labeled protein (magenta) at the working conditions (150 mM NaCl, 23°C, 10mg/mL glycinin). The condensate is only in contact with the fluid phase.

**Movie S2:** Confocal microscopy z-stack of a giant vesicle composed of DOPC:DPPC:Cholesterol 1:1:1 (green) displaying liquid-disordered/liquid-ordered phase coexistence in contact with a glycinin (magenta) at the working conditions. The condensate is only in contact with the liquid-disordered phase.

**Movie S3:** Confocal microscopy z-stack of a giant vesicle composed of DOPC:SM:Cholesterol 1:1:1 (green) displaying liquid-disordered/liquid-ordered phase coexistence in contact with a glycinin condensate (magenta) at the working conditions. The condensate is only in contact with the liquid-disordered phase.

**Movie S4:** Confocal microscopy z-stack of a giant vesicle composed of DOPC (green) in contact with a glycinin condensate (magenta) at the working conditions. Tubes are formed at the membrane-condensate interface protruding into the condensate phase.

**Movie S5:** Confocal microscopy z-stack of a giant vesicle composed of DOPC:DPPC 1:1 (green) in contact with a glycinin condensate (magenta) at the working conditions. Tubes are formed at the membrane-condensate interface protruding into the condensate phase.

**Movie S6:** STED microscopy z-stack of the membrane channel of the condensate-membrane interface for a giant vesicle composed of DOPC in contact with a glycinin condensate at the working conditions. Tubes formed at the interface are resolved, see Figure 5D for tube diameter values.

**Movie S7:** STED microscopy z-stack of the membrane channel of the condensate-membrane interface for a giant vesicle composed of DOPC:DPPC 1:1 in contact with a glycinin condensate (magenta) at the working conditions. Tubes formed at the interface are resolved, see Figure 5D for tube diameter values.

**Movie S8:** Confocal microscopy z-stack of the membrane channel (green) of a giant vesicle composed of DOPC in contact with a glycinin solution at 365 mM NaCl. Nanotubes are adhered to the outer membrane surface and a double-membrane sheet is observed at one of the vesicle poles.
